# The Shu Complex Prevents Mutagenesis and Cytotoxicity of Single-Strand Specific Alkylation Lesions

**DOI:** 10.1101/2021.04.09.439200

**Authors:** Braulio Bonilla, Alexander J. Brown, Sarah R. Hengel, Kyle S. Rapchak, Debra Mitchell, Catherine A. Pressimone, Adeola A. Fagunloye, Thong T. Luong, Hani S. Zaher, Nima Mosammaparast, Ewa P. Malc, Piotr A. Mieczkowski, Steven A. Roberts, Kara A. Bernstein

## Abstract

Three-methyl cytosine (3meC) are toxic DNA lesions, blocking base pairing. Bacteria and humans, express members of the AlkB enzymes family, which directly remove 3meC. However, other organisms, including budding yeast, lack this class of enzymes. It remains an unanswered evolutionary question as to how yeast repairs 3meC, particularly in single-stranded DNA. The yeast Shu complex, a conserved homologous recombination factor, aids in preventing replication-associated mutagenesis from DNA base damaging agents such as methyl methanesulfonate (MMS). We found that MMS-treated Shu complex-deficient cells, exhibit a genome-wide increase in A:T and G:C substitutions mutations. The G:C substitutions displayed transcriptional and replicational asymmetries consistent with mutations resulting from 3meC. Ectopic expression of a human AlkB homolog in Shu-deficient yeast rescues MMS-induced growth defects and increased mutagenesis. Finally, the Shu complex exhibits increased affinity for 3meC-containing DNA. Thus, our work identifies a novel mechanism for coping with alkylation adducts.

## INTRODUCTION

Alkylating agents such as methyl methanesulfonate (MMS) induce a diverse set of base lesions that are recognized and repaired by the base excision repair (BER) pathway or direct repair enzymes such as the AlkB family (Fu, Calvo et al., 2012, Yi & He, 2013). However, when these lesions, or its repair intermediates, are encountered by a replisome, replication fork stalling can occur (Shrivastav, Li et al., 2010, Sobol, Kartalou et al., 2003). In this scenario, DNA base damage is preferentially bypassed using homologous recombination (HR) or translesion synthesis (TLS), postponing its repair but allowing replication to be completed in a timely fashion. These pathways are often referred to as post-replication repair (PRR) or DNA damage tolerance (DDT) and are best described in the budding yeast *Saccharomyces cerevisiae* (Arbel, Liefshitz et al., 2020, Boiteux & Jinks-Robertson, 2013, Ulrich, 2007).

In yeast, HR-mediated PRR is an error-free pathway and is dependent on the polyubiquitination of PCNA by the Mms2-Rad5-Ubc13 complex (Arbel et al., 2020). Lesion bypass is achieved by Rad51 filament formation, and recombination between sister chromatids to fill the single-stranded DNA (ssDNA) gaps originated from the stalling of replicative polymerases. While HR can bypass these lesions in an error-free manner, TLS can also bypass this damage. However, TLS may lead to mutations and is often referred as error-prone lesion bypass. In yeast, TLS serves as an alternative pathway to error-free lesion bypass, as disruption of genes involved in the error-free PRR pathway leads to increased mutations that is dependent on TLS (Broomfield, Chow et al., 1998, Huang, Rio et al., 2003, Swanson, Morey et al., 1999).

The study of the error-free PRR pathway is challenging due to the diverse roles of HR proteins in additional processes, such as DSB repair. The Shu complex is an evolutionarily conserved HR factor that we recently discovered to function in strand-specific DNA damage tolerance during replication and thus provides an excellent model to specifically study HR proteins in the context of PRR (Godin, Meslin et al., 2015, Godin, Sullivan et al., 2016b, Martino & Bernstein, 2016, Rosenbaum, Bonilla et al., 2019). In *S. cerevisiae*, the Shu complex is a heterotetramer formed by the SWIM domain-containing protein, Shu2, and the Rad51 paralogs, Csm2, Psy3, and Shu1. The Shu complex promotes Rad51 filament formation, a key step for HR (Godin et al., 2016b). Consistent with their role in DSB repair, most HR genes deletions lead to increased sensitivity to DSB-inducing agents. However, this is not the case with the Shu complex, as its mutants are primarily sensitive to the alkylating agent MMS, but not to the DSB inducing agents such as ionizing radiation (Godin, Zhang et al., 2016c, Shor, Weinstein et al., 2005). This makes the Shu complex attractive to dissect the role of HR in the tolerance of replication-associated DNA damage.

Previous studies from our group and others demonstrated that the Shu complex operates in the error-free branch of the PRR to tolerate DNA damage from MMS-induced lesions (Ball, Zhang et al., 2009, Godin, Wier et al., 2013). Despite these findings, it has remained unknown which MMS-induced lesions, or repair intermediates, the Shu complex is important for. Our previous results uncovered genetic interactions between factors of the BER pathway and the Shu complex upon MMS treatment (Godin et al., 2016c). Notably, cells lacking the BER abasic (AP) endonucleases and AP lyases that process AP sites exhibit exquisite sensitivity towards alkylating agents and display a three order of magnitude increase in mutations rates when the Shu complex is disrupted. We recently demonstrated that the Shu complex is important for tolerance of AP sites (Rosenbaum et al., 2019). However, it remains outstanding question as to whether the Shu complex may recognize other MMS-induced fork blocking lesions.

Here, we addressed whether the Shu complex function is specific for AP sites or if it is important for recognition of a broader range of DNA lesions. To do this, we performed whole genome sequencing of Shu mutant cells (i.e. *csm2*Δ) that had been chronically exposed to MMS. In turn, this allowed us to obtain an unbiased spectrum of mutations the Shu complex helps to mitigate. We determined that the Shu complex prevents mutations at both A:T and C:G base pairs. However, transcriptional and replicative asymmetries in C:G mutations suggest a novel role for the Shu complex in the tolerance of 3meC, in addition to AP sites. Importantly, unlike bacteria and human cells that have an enzyme that directly repairs 3meC, this family of enzymes is absent in *S. cerevisiae* and therefore it has remained unknown how 3meC are repaired in yeast (Sedgwick, Bates et al., 2007). Our data suggest that the Shu complex is important for 3meC tolerance. Indeed, expression of human *ALKBH2*, responsible for 3meC repair, specifically rescues the MMS-sensitivity of Shu-complex mutant cells and alleviates their MMS-induced mutagenesis. In contrast to Shu complex mutants, we observe that *ALKBH2* expression very weakly rescues the MMS sensitivity associated with HR deletion strains. Finally, we find that the Shu complex DNA-binding subunits, Csm2-Psy3, have higher affinity for double-flap substrates containing 3meC relative to an unmodified substrate. Altogether, our findings reveal a previously unappreciated role for the Shu complex in mediating damage tolerance of 3meC in single-stranded DNA, finally uncovering how yeast tolerate these highly toxic lesions. Additionally, the ability of the Shu complex to facilitate tolerance of 3meC DNA damage through homology-directly bypass likely highlights an important repair pathway for these lesions even in organisms that express AlkB homologs.

## RESULTS

### Unbiased genome-wide analysis of mutation patterns suggests that the Shu complex function in the error-free bypass of specific MMS-induced lesions

To determine the identity of MMS-induced lesions that the Shu complex helps the replication fork to bypass, we chronically exposed Shu complex-deficient cells to MMS. In particular, wild-type (WT) or *csm2*Δ cells were plated on MMS-containing medium and then individual colonies were transferred every two days onto fresh medium containing 0.008% MMS, for a total of ten passages. We then extracted genomic DNA from these colonies and performed whole-genome sequencing to measure mutation frequencies. Consistent with previous findings, *csm2*Δ cells accumulated a median 12.7-fold more mutations than WT upon MMS treatment (**Figure 1A**) (Godin et al., 2013, Godin et al., 2016c). To infer which DNA lesions caused the mutations, we analyzed the substitution patterns considering the MMS-induced lesion profile (Beranek, 1990, Sikora, Mielecki et al., 2010, Wyatt & Pittman, 2006). 3meA is the most common MMS-induced lesion at A:T base pairs in dsDNA and can itself be mutagenic or converted to a mutagenic AP site by the Mag1 glycosylase or by spontaneous depurination (**Figure 1B** and **1C**) (Shrivastav et al., 2010). When these AP sites are encountered by a replicative polymerase, they can lead to its stalling. TLS activity on 3meA-derived AP sites leads to an A->G and A->T substitution pattern (**Figure 1C**) due to the tendency of Rev1 or the replicative polymerase δ incorporating a C or A on AP sites, respectively (**Figure 1C**) (Chan, Resnick et al., 2013, Haracska, Unk et al., 2001, Hoopes, Hughes et al., 2017). Consequently, most substitutions at A:T bases in MMS-treated WT yeast A to G and A to T (**Figure 1B**). Previous analyses of MMS-treated *mag1*Δ yeast revealed that deletion of the glycosylase responsible for initiating base excision repair of 3meA resulted in a mutation spectrum also predominantly composed of A to G and A to T substitutions (Mao, Brown et al., 2017), indicating that these two mutation types are likely directly induced by 3meA as opposed to 3meA-derived AP sites. MMS-treated *csm2*Δ cells (**Figure 1B**) displayed elevated A->G transitions and A->T transversions, consistent with a function for the Shu complex in the bypass of 3meA or 3meA-derived AP sites, as we recently demonstrated (Godin et al., 2016c, Rosenbaum et al., 2019).

**Figure 1.**
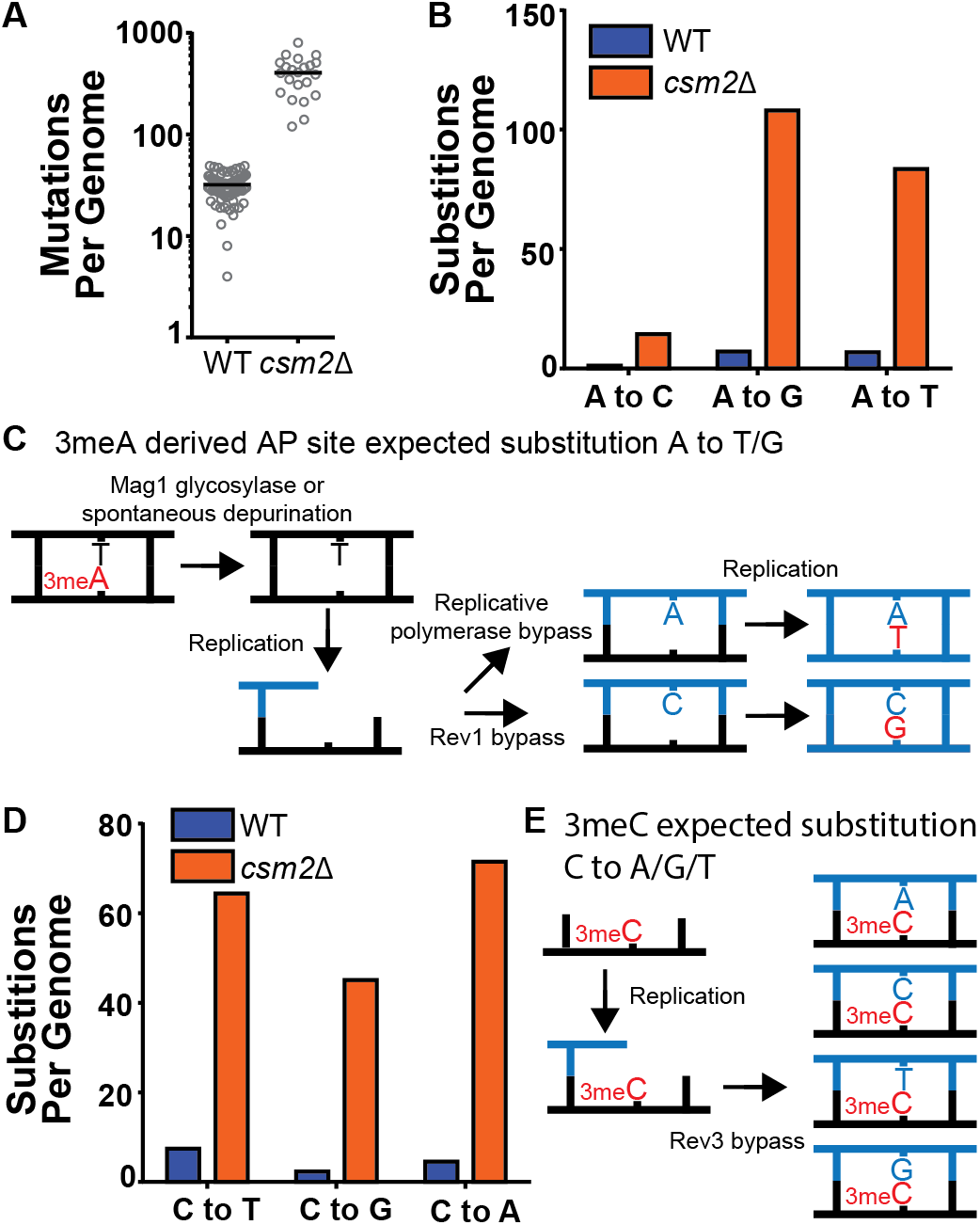
*csm2Δ* Cells Chronically Exposed to MMS Exhibit Substitution Patterns Consistent with TLS Activity Bypassing AP Sites and 3meC. WT and *csm2*Δ cells were chronically exposed to MMS. WT and *csm2*Δ cells were chronically exposed to 0.008% MMS by plating individual colonies onto rich medium containing MMS, after 2 days of growth, the colonies were platted onto fresh medium containing MMS for 10 passages. DNA was extracted from 90 colonies per genotype and deep sequenced. (A) The number of mutations per genome for WT or *csm2*Δ cells chronically MMS-exposed. Horizontal bar indicates the median value for each genotype. (B) The average number of each type of A:T substitution per genome in MMS-treated WT and *csm2*Δ cells. (C) Schematic of how 3meA derived AP sites result in A to T/G mutations. 3meA is removed by the Mag1 glycosylase or spontaneously depurinated resulting in an AP site. During DNA replication, the replicative polymerase bypasses the AP site resulting in a T mutation or alternatively Rev1 bypasses the AP site resulting in a G mutation. (D) The average number of each type of G:C substitution per genome in MMS-treated WT and *csm2*Δ cells. (E) Schematic of how 3meC results in C to A/G/T base substitutions. 3meC occurs primarily in ssDNA and during replication, Rev3 mediated bypass results in incorporation of A, T, G, or C nucleotides.

7meG is the most common MMS-induced DNA adduct at G:C base pairs and is not itself mutagenic (Shrivastav et al., 2010). However, it is eliminated by spontaneous depurination, or Mag1 excision, which both lead to AP sites (Bjoras, Klungland et al., 1995, Shrivastav et al., 2010, Wyatt, Allan et al., 1999). When AP sites are generated from guanines, their bypass by Rev1 is error-free because it incorporates a C across the missing base. Alternatively, bypass by the replicative polymerase results in the incorporation of A across the missing base, which leads to a G to T transversion (**Figure S1**). Interestingly, the MMS-induced mutation profile at G:C base pairs in WT yeast, not only includes G->T mutations but also high levels of G->A and G->C mutations (**Figure 1D**), suggesting that 7meG-derived AP sites are unlikely to cause these mutations. Deletion of *CSM2* results in a similar spectrum of MMS-induced G:C substitutions compared to that observed in WT yeast, but at elevated frequency, indicating that the Shu complex mediates tolerance of the same MMS-induced lesion that contributes to G:C base pair mutagenesis in WT cells, possibly N3-methylcytosine (3meC). 3meC is a mutagenic DNA adduct predominantly induced by MMS at ssDNA and can block the replicative polymerase (Shrivastav et al., 2010). Rev3-mediated TLS bypass of 3meC leads to the incorporation of a random nucleotide across the lesion, leading to a mutational pattern consistent with the one we observed in WT and *csm2*Δ yeast (**Figure 1E**) (Saini, Sterling et al., 2020, Yang, Gordenin et al., 2010), indicating that the Shu complex may also bypass this ssDNA specific alkylation lesion.

DNA lesions that arise at ssDNA often result in mutation patterns that exhibit strand bias near origins of replication and at highly transcribed genes (Roberts, Sterling et al., 2012, Saini & Gordenin, 2020). Additionally, lesions subject to preferential repair of the transcribed DNA strand by transcription-coupled nucleotide excision repair (TC-NER) can also display transcriptional asymmetry (i.e. having higher mutation densities on the non-transcribed strand compared to the transcribed stand). We therefore evaluated whether mutations from our whole genome sequenced yeast displayed either transcriptional or replicative asymmetry (**Figure 2**). A:T mutations in both MMS-treated WT and *csm2Δ* yeast showed very little replicative asymmetry, but displayed transcriptional asymmetry, similar to that seen in MMS-treated *mag1*Δ yeast (Mao et al., 2017). The transcriptional asymmetry of A:T mutations in MMS-treated *mag1*Δ yeast results from TC-NER activity on unrepaired 3meA, indicating that the Shu complex may help prevent mutations associated with 3meA (Beranek, 1990, Sikora et al., 2010, Wyatt & Pittman, 2006). However, increased mutagenesis of the minor ssDNA specific lesion N1-methyl adenine (1meA) would also be expected to produce higher mutation densities on the non-transcribed strand. Therefore, we cannot exclude the possibility that 1meA contributes to the increased MMS-induced mutation rate in *csm2Δ* cells. G:C mutations in MMS-treated WT yeast displayed little to no transcriptional or replicative strand asymmetry, while the same mutations in MMS-exposed *csm2*Δ cells exhibited both types of strand bias (**Figure 2**). The transcriptional asymmetry displayed higher densities of C mutations on the non-transcribed strand, consistent with either TC-NER removal of 3meC or preferential formation of 3meCs on the transiently ssDNA non-transcribed strand, instead of an asymmetry derived for TC-NER of 7meG. We additionally note that transcriptional asymmetry of the 3meC in MMS-treated *csm2*Δ extends up to 167 bp upstream of the transcriptional start site. This may suggest that either TC-NER can function within this region or some ssDNA is assessable on the non-transcribed strand due to the binding of TFIIH to the promoter region upstream of the TSS. Likewise, replicative asymmetry indicated elevated C mutations associated with the lagging strand template across the yeast genome, suggesting that 3meC lesions formed during lagging strand synthesis are bypassed in an error-free manner in Shu proficient yeast.

**Figure 2.**
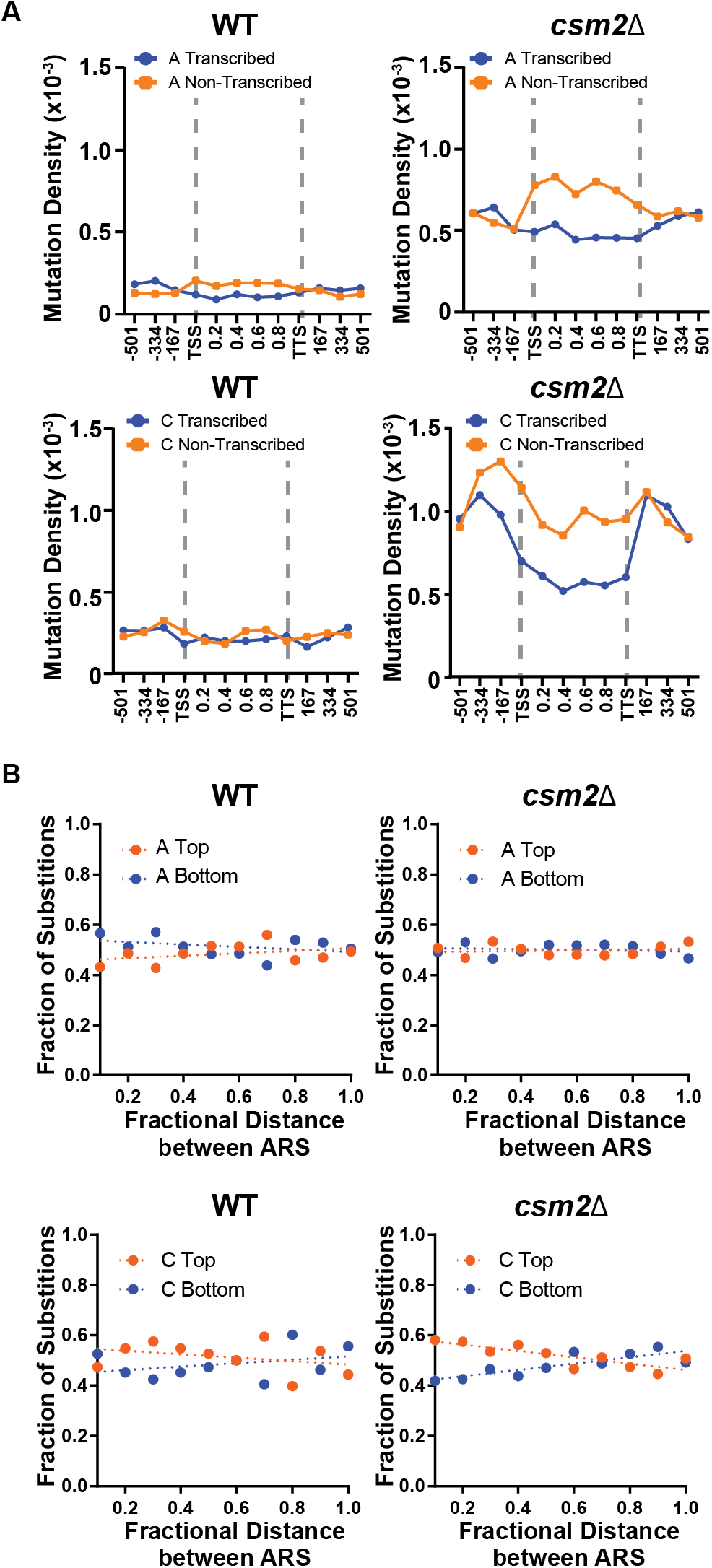
Transcriptional and Replicative Strand Biases of MMS-Induced Substitutions. (A) The density of A mutations (i.e. the fraction of A bases mutated) or C mutations on the transcribed (blue) and non-transcribed (orange) strand across yeast transcripts in WT and *csm2*Δ cells. Transcript regions were broken into fractional bins of 0.2 of the transcript length and the density of A or C mutations determined per bin. 3 additional bins of 500 bp each where also included upstream of the transcription start site (TSS) and downstream of the transcription termination site (TTS). (B) The fraction of A and C mutations occurring on the top (orange) and bottom (blue) strands across replication units in the genomes of WT and *csm2*Δ cells. Replication units were broken into 0.1 fractional bins between neighboring origins of replication and the fraction of A mutations or C mutations associated with each strand were calculated per bin. The fraction of mutations for each strand across the replication unit was fitted with linear regression lines (dashed lines).

### *ALKBH2* expression rescues the MMS-induced phenotypes of Shu complex deficient cells

Based on the analysis of the mutation patterns, we hypothesized that the Shu complex promotes the error-free bypass of 3meC. 3meC can be repaired by the AlkB family of Fe(II)/α-Ketoglutarate-dependent dioxygenases (Fedeles, Singh et al., 2015). This family is conserved from bacteria to humans; however, no yeast homolog has been found. Therefore, as of yet it remains unknown how yeast tolerates and repairs the highly toxic 3meC. In humans, there are nine AlkB homologs with ALKBH2 and ALKBH3 are responsible for 3meC DNA repair (Fedeles et al., 2015, Yi & He, 2013). We, therefore, reasoned that ectopic expression of AlkB homologs would rescue the MMS-induced phenotypes observed in Shu complex deficient cells. To test this, we took advantage of the lack of an AlkB homolog in yeast by ectopically expressing human AlkB homologs, *ALKBH2*, or *ALKBH3*, in *csm2*Δ cells and analyzed their effect on MMS sensitivity. *ALKBH2* and *ALKBH3* were expressed under the constitutive GAP promoter using a low-copy CEN plasmid. We find that both *ALKBH2* and *ALKBH3* expression leads to a partial rescue of the growth defects observed in MMS-exposed *csm2Δ* cells, with *ALKBH2* showing a stronger rescue (**Figure 3A**). Therefore, we focused on *ALKBH2* for the remainder of the experiments. As expected, *ALKBH2* expression very mildly rescues the MMS sensitivity of WT cells (**Figure 3A**). The growth defect rescue observed in MMS-treated *csm2Δ* cells depends on ALKBH2’s enzymatic activity since expression of an *ALKBH2* <u>c</u>atalytic <u>d</u>ead mutant (ALKBH2-V101R,F120E herein referred to as alkbh2-CD) (Monsen, Sundheim et al., 2010) does not rescue *csm2*Δ cell viability (**Figure 3B**). We confirmed that the inability of alkbh2-CD to rescue *csm2*Δ cells was not due to lower protein expression (**Figure S2)**. These findings are not specific to *csm2*Δ as *ALKBH2* rescues the MMS sensitivity of the other Shu complex members to the same extent (**Figure 3C**). The effect of *ALKBH2* expression on *csm2*Δ cell viability is specific for MMS damage as *ALKBH2* expression does not rescue the known growth defect observed in UV-treated *csm2Δ rev3Δ* double mutant cells (Xu, Ball et al., 2013) (**Figure 3D**).

**Figure 3.**
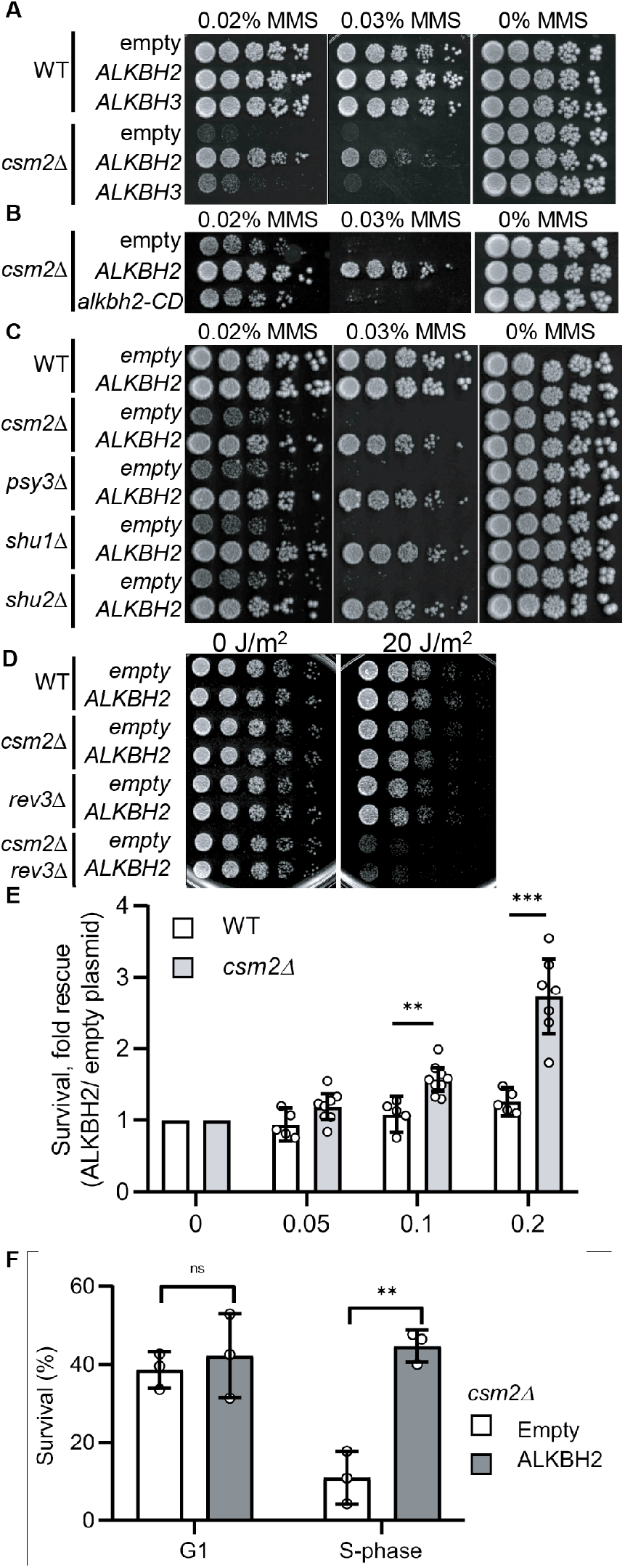
Expression of Human *ALKBH2* Rescues the MMS Sensitivity of *csm2Δ* Cells. (A) *csm2Δ* cells expressing *ALKBH2* exhibit decreased MMS sensitivity. Five-fold serial dilution of WT or *csm2*Δ cells transformed with an empty plasmid, a plasmid expressing *ALKBH2* or a plasmid expressing *ALKBH3* onto rich YPD medium or YPD medium containing the indicated MMS concentration were incubated for 2 days at 30 °C prior to being photographed. (B) The enzymatic activity of ALKBH2 is required for the rescue of the MMS sensitivity of *csm2*Δ cells. *csm2*Δ cells transformed with an empty plasmid, a plasmid expressing *ALKBH2* or a plasmid expressing a catalytic dead *ALKBH2* mutant were diluted and plated as described in (A) and incubated for three days at 30°C prior to being photographed. (C) Expression of *ALKBH2* does not rescue the increased UV sensitivity observed in *csm2Δ rev3Δ* double mutants. Five-fold serial dilution of WT, *csm2Δ*, *rev3Δ*, or *rev3Δ csm2Δ* cells were transformed with an empty plasmid or a plasmid expressing *ALKBH2* and five-fold serially diluted onto rich YPD or rich YPD medium exposed to 20 J/m^2^ ultra-violet (UV), and incubated for 2 days at 30°C prior to being photographed. An untreated plate (0 J/m^2^) serves as a loading control. (D) *ALKBH2* expression rescues the MMS sensitivity of cells with deletions of the four Shu complex genes. WT, *csm2*Δ, *psy3*Δ, *shu1*Δ, or *shu2*Δ cells transformed with an empty plasmid, or a plasmid expressing *ALKBH2* were five-fold serially diluted, plated and analyzed as described in (B). (E) *csm2Δ* cells expressing *ALKBH2* exhibit increased survival after acute MMS treatment. YPD liquid cultures of WT or *csm2*Δ cells transformed with an empty plasmid or a plasmid expressing *ALKBH2* were treated with the indicated concentration of MMS following plating onto rich YPD medium. Colony number was assessed after incubation for two days at 30°C. Fold rescue of cellular survival represents the ratio of the survival of cells expressing *ALKBH2* relative to the survival of cells expressing the empty plasmid. Survival represents the number of colonies as a percentage of the colonies obtained without MMS treatment. The individual and mean values from five to nine experiments were plotted. Error bars indicate 95% confidence intervals. The *p*-values between WT and *csm2*Δ cells treated with 0.1% MMS and 0.2% MMS were calculated using an unpaired two-tailed Student’s t-test and were p ≤ 0.01 and p ≤ 0.001, respectively. (F) S-phase *csm2*Δ cells expressing *ALKBH2* exhibit increased survival after acute MMS treatment. WT or *csm2*Δ cells were synchronized on G1 with alpha factor and either released from G1 arrest or maintained in G1 in the presence or absence of 0.1% MMS. Cells were plated after 30 minutes of treatment and the colony number was assessed after incubation for two days at 30°C. Survival is calculated as described in (E). The mean values from three experiments were plotted with standard deviations. The *p*-values between control (empty plasmid) and *ALKBH2* expressing cells were calculated using an unpaired two-tailed Student’s t-test and were p>0.05 (n.s.) and p ≤ 0.001 for the G1 and S-phase cells, respectively.

Next, we analyzed the effect of *ALKBH2* expression on *csm2Δ* cells acutely exposed to MMS. To do this, we treated *ALKBH2*-expressing WT or *csm2Δ* cultures with 0.1% MMS for 30 minutes. We then assessed cell survival by counting viable colonies after two days of growth. We observe that *ALKBH2* expression leads to a dose-dependent increase in the survival of *csm2Δ* cells, whereas in WT cells, survival is only mildly rescued (**Figure 3E**).

Since 3meC only occurs at ssDNA, which is a physiological intermediate of DNA replication, we reasoned that *ALKBH2* expression would preferentially rescue the survival of cells that are progressing through S-phase and therefore more vulnerable to 3meC-induced toxicity. To test this, we compared the survival of *ALKBH2*-expressing *csm2Δ* cells arrested in G1 in MMS containing media with *ALKBH2*-expressing *csm2Δ* cells progressing through S-phase. In agreement with our rationale, *ALKBH2* was found to rescue the survival phenotype only for cells progressing in the S phase of cell cycle (**Figure 3F**). This is consistent with previous data in human cells suggesting that *ALKBH2* functions preferentially in S-phase (Gilljam, Feyzi et al., 2009).

### *ALKBH2* expression alleviates the MMS-induced mutations observed in Shu complex mutant, *csm2*Δ

Since 3meC is a mutagenic lesion, we asked whether *ALKBH2* expression would alleviate the observed mutational load of MMS-exposed *csm2Δ* cells. To do this, we utilized the *CAN1* reporter assay. The *CAN1* gene encodes for an arginine permease, therefore when cells are exposed to the toxic arginine analog canavanine, only cells that acquire mutations in the *CAN1* gene grow. *ALKBH2* expression decreased the mutation rates in MMS-exposed *csm2Δ* cells (**Figure 4A**). This decreased mutation rate is specific for alkylation lesions as *ALKBH2* expression fails to rescue the increase in spontaneous mutations observed in a Shu complex mutant, *csm2Δ* cells (**Figure 4A**). Furthermore, the rescue of *csm2*Δ mutation rate by *ALKBH2* depends on *ALKBH2* catalytic activity (**Figure S3**). These spontaneous mutations are likely due to the TLS-mediated bypass of AP sites (Ball et al., 2009, Godin et al., 2016c, Rosenbaum et al., 2019, Shor et al., 2005, Shrivastav et al., 2010). To determine whether the decrease in mutations in MMS-treated, *ALKBH2*-expressing *csm2Δ* cells corresponds to the repair of 3meC, we sequenced the *CAN1* gene of canavanine resistant clones. Can-R isolates from MMS-treated *csm2Δ* cells displayed frequent mutation of the *CAN1* gene with substitutions at A:T and C:G bases as well as single nucleotide deletions and complex events consisting of two mutations separated by less than 10 bp that are a signature of error-prone trans-lesion synthesis (Northam et al., 2010, Sakamoto et al., 2007). Accordingly, *ALKBH2* expression in *csm2Δ* cells decreased the frequency of deletions, complex mutations, and substitutions at C:G base pairs (**Figure 4B**). These findings support a model in which the Shu complex prevents the cytotoxicity and mutagenicity of 3meC and in its absence, yeast are reliant on TLS to bypass these lesions resulting an increase of C:G substitutions, small deletions, and complex mutations.

**Figure 4.**
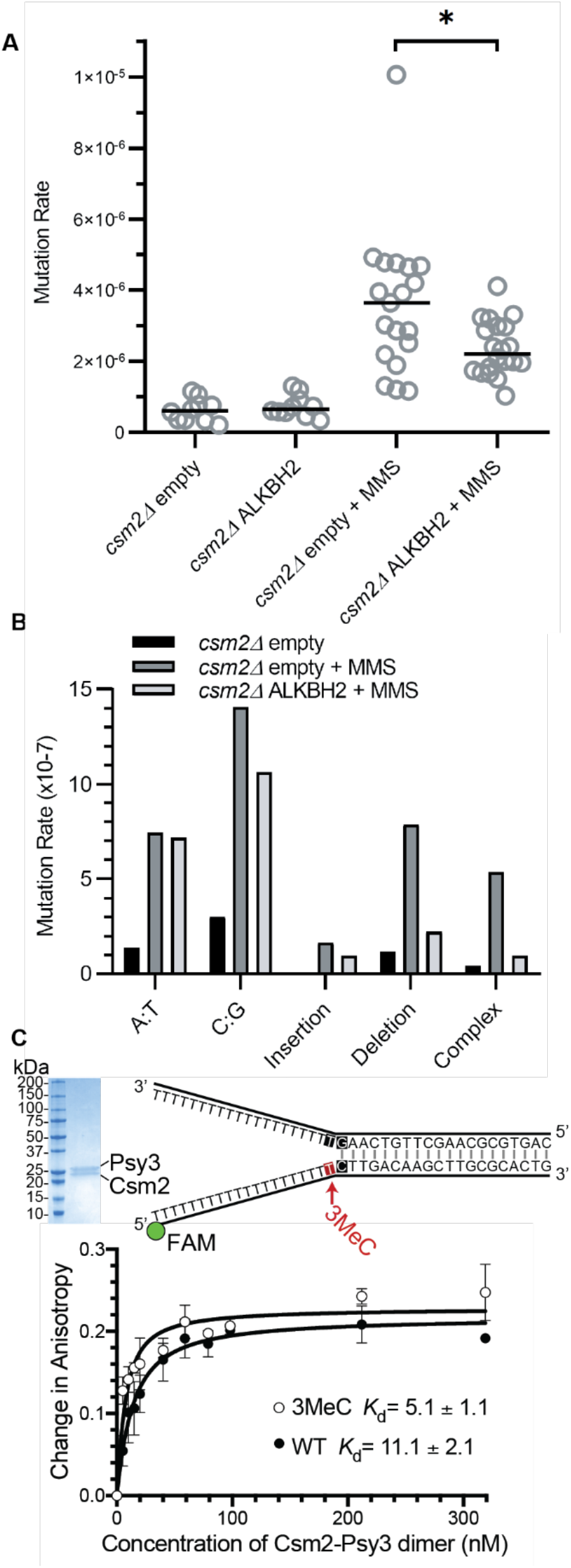
*ALKBH2* Expression Rescues the 3meC-Induced Mutagenesis Observed in MMS Exposed *csm2Δ* Cells. (A) *csm2Δ* cells expressing *ALKBH2* exhibit reduced MMS-induced mutation rate. Spontaneous and MMS-induced mutation rates at the *CAN1* locus were measured in *csm2*Δ cells transformed with an empty plasmid or a plasmid expressing *ALKBH2*. Each measurement (dots) and the median value of 10 to 20 experiments (horizontal bar) were plotted. The *p*-values between control (empty plasmid) and *ALKBH2* expressing cells were calculated using a Mann-Whitney Ranked sum test and were p>0.05 and p < 0.03 for the for the untreated and MMS treated samples respectively. (B) Sequencing of the *CAN1* gene in canavanine resistant colonies was used to calculate the frequency of MMS-induced substitutions, insertions, deletions, and complex mutations in *csm2*Δ cells transformed with either an empty vector or *ALKBH2* expression vector. ALKBH2 expression significantly alters the MMS-induced mutation spectra (p = 0.018 by Chi-square analysis comparing the number of A:T substitutions, C:G substitutions, insertions, deletions, and complex mutations between MMS-treated *csm2*Δ cells containing an empty vector or *ALKBH2* expression vector). The spectrum contains a reduction in C:G substitutions, deletions, and complex mutations, while A:T substitutions and insertions are unchanged. (C) The Csm2-Psy3 protein has high affinity for a double-flap DNA substrate containing a 3meC lesion. A Coomassie stained SDS-PAGE gel of recombinant Csm2-Psy3, which run at 27.7 kDa and 28 kDa, respectively. Equilibrium titrations were performed by titrating Csm2-Psy3 into the FAM-labeled double-flap substrate in the presence (4.53 nM 3meC) or absence (5.11 nM WT) of a 3meC at the indicated position and anisotropy was measured. Triplicate experiments were performed with standard deviations plotted. The data were fit to a quadratic equation (one-site binding model assumed) and dissociation constants (*K_d_*) were calculated. Note the improved binding compared to previous studies (Rosenbaum et al., 2019) due to increased protein activity from improved constructs.

We previously demonstrated that the Shu complex preferentially binds to double-flap DNA substrates and has a two-fold higher affinity for double-flap DNA containing an AP site analog (Godin et al., 2016c, Rosenbaum et al., 2019). Therefore, we asked whether the DNA binding subunits of the Shu complex, Csm2-Psy3, would similarly have higher affinity for double-flap DNA containing 3MeC. To address this, we examined the affinity of recombinant Csm2-Psy3 to a FAM-labeled double-flap substrate in the presence or absence of 3meC by fluorescence anisotropy (**Figure 4C**). Indeed, Csm2-Psy3 has a two-fold improved affinity for a double-flap substrate containing a 3meC lesion on the ssDNA at the ssDNA/dsDNA junction relative to unmodified substrate [3meC, *K*_d_= 5.1 ± 1.1; C, *K*_d_= 11.1 ± 2.1; **Figure 4C**]. These results suggest that the Shu complex increases error-free bypass of 3meC by preferentially recognizing 3meC lesions on ssDNA in double-flap substrates.

### 3meC repair in yeast is channeled through error-free post-replicative repair

The Shu complex directly functions with the canonical Rad51 paralogs, Rad55-Rad57, and Rad52 to promote HR through Rad51 filament formation (Gaines, Godin et al., 2015, Godin et al., 2013). In the context of error-free post-replicative repair, the Shu complex functions downstream of poly-ubiquitination of PCNA by Rad5-Ubc13-Mms2 (Xu et al., 2013). Therefore, we asked whether the rescue of the MMS-induced phenotypes by *ALKBH2* depends on Shu complex function as a Rad51 mediator. To address this question, we utilized a Csm2 mutant, *csm2-F46A*, that cannot stimulate Rad51 filament formation due to loss of its protein interaction with Rad55-Rad57 (Gaines et al., 2015, Godin et al., 2013). Suggesting that the Shu complex mediator function enables bypass of 3meC, we find that *ALKBH2* expression suppresses the MMS sensitivity of a *csm2-F46A* mutant to the same extent as a *csm2*Δ cell (**Figure 5A**). Next, we asked whether the rescue of MMS sensitivity by *ALKBH2* would be observed in other HR mutants. To accomplish this, we ectopically expressed *ALKBH2* in cells with deletions of *CSM2, RAD51, RAD52, RAD55*, and *UBC13* and performed serial dilutions upon increasing-MMS dosage (**Figure 5B**). Surprisingly, *rad51*Δ, *rad52*Δ, and *rad55*Δ MMS sensitivity is not rescued by *ALKBH2* expression to the same extent as *csm2*Δ cells. In contrast, *ubc13*Δ MMS sensitivity is largely rescued by *ALKBH2* expression. This striking result suggests that the Shu complex functions primarily within Ubc13-initiated post-replicative repair pathway, while the canonical HR genes are also critical for the repair of DSBs induced by MMS through clustered lesions. Unlike Shu complex mutant cells, deletion of the canonical HR genes leaves cells vulnerable to toxic MMS-induced DSBs formed from lesions that are not subject to *ALKBH2* direct reversal.

**Figure 5.**
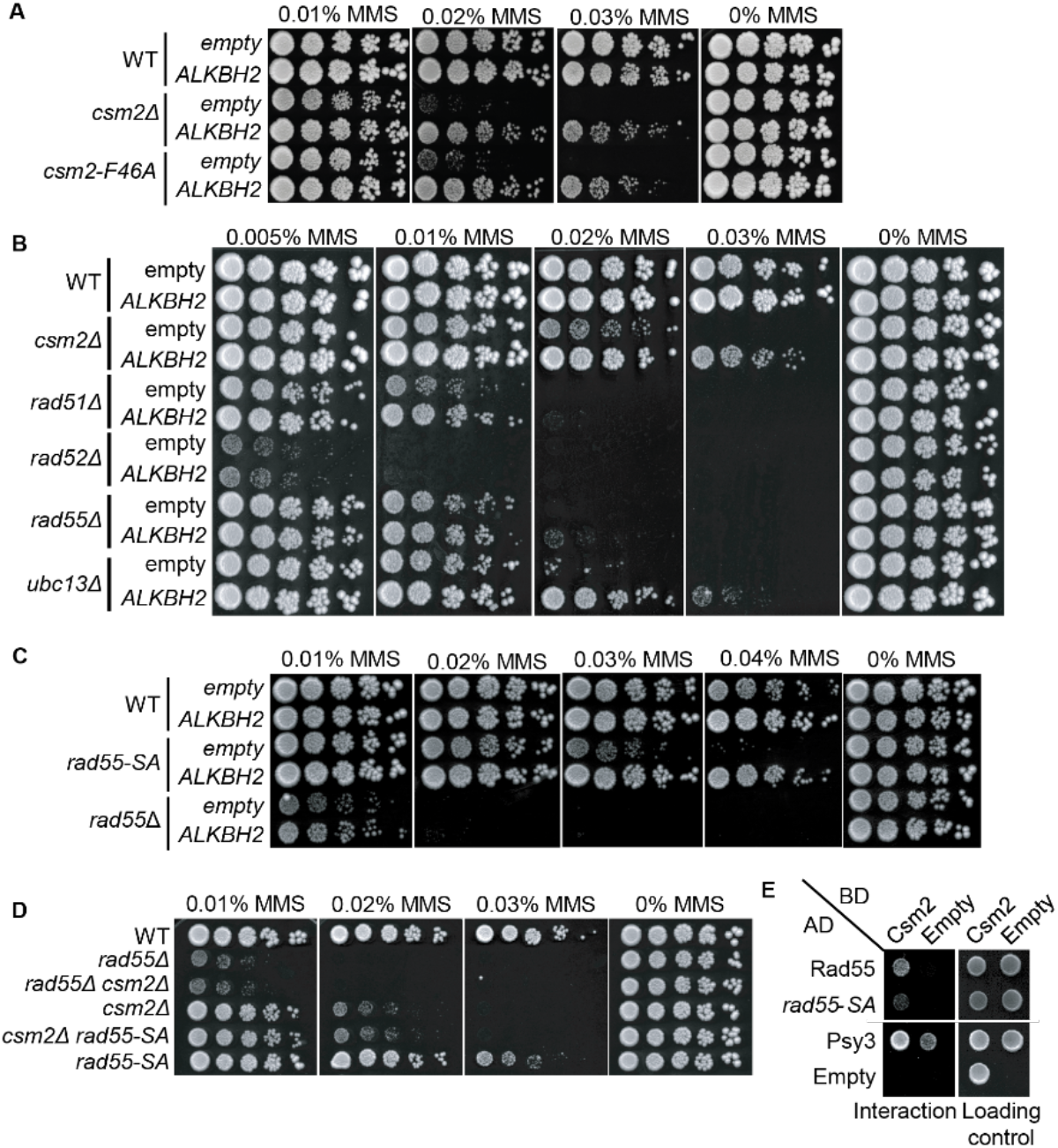
3meC are Bypassed by the Error-Free Post Replicative Repair (PRR) Pathway. (A) *ALKBH2* rescues the MMS sensitivity of a *csm2-F46A* mutant, which is deficient for its Rad51 mediator activity. Five-fold serial dilution of WT, *csm2Δ*, or *csm2-F46A* cells were transformed with an empty plasmid or a plasmid expressing *ALKBH2* onto rich YPD medium or YPD medium containing the indicated MMS concentration and incubated for 3 days at 30°C prior to being photographed. (B) Unlike PRR mutant *UBC13*, expression of *ALKBH2* mildly rescues the MMS sensitivity of HR factors, *RAD51, RAD52*, and *RAD55*. Five-fold serial dilution of WT, *csm2Δ*, *rad51Δ*, *rad52Δ*, *rad55Δ*, or *ubc13Δ* transformed with an empty plasmid or a plasmid expressing *ALKBH2* were five-fold serially diluted, plated, and analyzed as described in (A). (C) *rad55-S2,8,14A* cells expressing *ALKBH2* exhibit decreased MMS sensitivity. WT, *rad55-S2,8,14A*, or *rad55Δ* cells transformed with an empty plasmid or a plasmid expressing *ALKBH2* were five-fold serially diluted, plated, and analyzed as described in (A). (D) *csm2Δ* is epistatic to *rad55-S2,8,14A* for MMS damage. Cells with the indicated genotypes were five-fold serially diluted and plated as described in (A), and incubated for 2 days at 30°C prior to being photographed. (E) rad55-S2,8,14A exhibit an impaired yeast-2-hybrid (Y2H) interaction with Csm2. Y2H analysis of pGAD-*RAD55*, *rad55-S2,8,14A, PSY3*, or pGAD-C1 (Empty) with pGBD-*RAD57*, *CSM2*, pGBD-C1 (Empty). A Y2H interaction is indicated by plating equal cell numbers on SC medium lacking histidine, tryptophan, and leucine. Equal cell loading is determined by plating on SC medium lacking tryptophan and leucine used to select for the pGAD (AD) and pGBD (BD) plasmids.

To explore this idea further, we investigated the effect of *ALKBH2* in a *RAD55* phosphorylation mutant that is MMS sensitive while maintaining its DSB-repair proficiency (Herzberg, Bashkirov et al., 2006). In this *RAD55* mutant, three serine residues (2,8,14) are mutated to alanine residues (*rad55-S2,8,14A*). Since the *rad55-S2,8,14A* mutant cells largely phenocopy the defects observed in a Shu complex mutant (Herzberg et al., 2006), we asked whether Rad55 function in MMS-induced DNA damage may be uncoupled from its role in canonical DSB repair. Unlike *rad55*Δ cells, the *rad55-S2,8,14A* mutant MMS sensitivity is largely rescued by *ALKBH2* expression (**Figure 5C**). To further investigate the genetic relationship between the Shu complex and Rad55, we combined either *rad55*Δ or *rad55-S2,8,14A* with a *csm2*Δ mutant. As previously reported, *csm2*Δ *rad55*Δ double mutants exhibit the same MMS sensitivity as a *rad55*Δ mutant cell (Godin et al., 2013). In contrast, a *csm2*Δ *rad55-S2,8,14A* double mutant exhibits the same MMS sensitivity as a single *csm2*Δ cell (**Figure 5D**). This observation is surprising since the Shu complex is thought to function downstream of Rad55. However, it is consistent with the specificity of the Shu complex in directly recognizing and enabling tolerance of MMS-induced DNA lesions (Rosenbaum et al., 2019).

Rad55 directly interacts with Csm2-Psy3 (Gaines et al., 2015, Godin et al., 2013). While Rad55-S2,8,14A mutant maintains its protein interactions with Rad57, Rad51, and Rad52, its interaction with the Shu complex has yet to be determined (Herzberg et al., 2006). Therefore, the Shu complex may help recruit Rad55 to specific MMS-induced DNA lesions through an interaction with phosphorylated Rad55 or at the interface where Rad55 is phosphorylated. To test this, we performed Y2H analysis of Rad55 or Rad55-S2,8,14S with Csm2 (**Figure 5E**). Interestingly, we observe a reduced interaction between Rad55-S2,8,14S with Csm2 (**Figure 5E**). These results suggest that Rad55 phosphorylation may stimulate its interaction with Csm2 or that Csm2 interacts with Rad55 in that region. In the context of MMS-induced DNA damage, our findings suggest that the Shu complex is likely contributing to the recognition of specific MMS-induced lesions and recruiting the HR machinery, such as Rad55-Rad57, to the lesion to facilitate their bypass and enable their repair following replication. Furthermore, other factors, such as a stalled replication fork, may also contribute to their recruitment.

## DISCUSSION

Among the many alkylation-induced DNA lesions, 3meC is well noted due to its cytotoxic and mutagenic impact during DNA replication (Nieminuszczy, Mielecki et al., 2009, Shrivastav et al., 2010, Sikora et al., 2010). In eukaryotes, the adduct is formed endogenously from S-adenosyl methionine (SAM) reactivity with DNA and from the enzymatic activity of DNA methyltransferases, and exogenously from alkylating agents such as nitrosamines, which are present in the tobacco smoke, temozolomide, or MMS (Chatterjee & Walker, 2017, Dango, Mosammaparast et al., 2011, Pataillot-Meakin, Pillay et al., 2016, Rosic, Amouroux et al., 2018). 3meC occurs primarily in ssDNA and can stall the replicative polymerases. Unlike bacteria and higher eukaryotes, yeast does not encode for an enzyme capable of directly repairing 3meC from ssDNA (Admiraal, Eyler et al., 2019, Sedgwick et al., 2007). Therefore, yeast rely on bypass mechanisms to complete DNA replication. Here, we utilized the budding yeast model to demonstrate that the Shu complex facilitates an HR-mediated error-free bypass of alkylation damage, including 3meC and AP sites, to prevent mutagenesis and toxicity.

The Shu complex is primarily involved in tolerance of replicative base-template damage, being dispensable for DSB repair (Ball et al., 2009, Godin et al., 2016c). This makes the Shu complex ideal to dissect the role of HR during bypass of specific base lesions. The major phenotypes observed in MMS-exposed Shu complex disrupted cells are decreased cell survival and elevated mutation frequency (Ball et al., 2009, Godin et al., 2016b, Shor et al., 2005). We observe that *ALKBH2* expression specifically alleviates the MMS-induced phenotypes of Shu complex disrupted cells (**Figures 3 and 4**). This *ALKBH2* rescue is partial, which is consistent with the Shu complex role in tolerance of another MMS-induced lesion, an AP site (Godin, Lee et al., 2016a, Rosenbaum et al., 2019). Although it is not possible to specifically induce 3meC, it is possible to regulate the occurrence of its template, by controlling the amount of ssDNA in MMS-exposed cells. It would be interesting to observe how the rescue by expression of *ALKBH2* correlates with the amount of ssDNA. Consistently, our results show that *ALKBH2* is only able to rescue the MMS sensitivity of Shu complex mutant cells that are progressing through S-phase and therefore exhibit ssDNA intermediates (**Figure 3E**). Furthermore, we find that Shu complex members, Csm2-Psy3, have a higher binding affinity for a double-flap substrate containing 3meC lesions on the ssDNA lagging strand (**Figure 4C**).

The error-free PRR pathway requires polyubiquination of PCNA by Mms2-Ubc13-Rad5 and the activity of the core HR machinery (Branzei & Szakal, 2017, Xu, Blackwell et al., 2015). Interestingly, the deletion of other HR factors, such as *RAD51*, are only partially rescued by *ALKBH2* expression (**Figure 5B**). This can be explained by their critical role in DSB repair, an activity for which the Shu complex is largely dispensable. Our results are consistent with 3meC being bypassed by the HR branch of the PRR pathway (**Figure 6**) and the notion that the Shu complex is an HR factor specialized for replication-damage.

**Figure 6.**
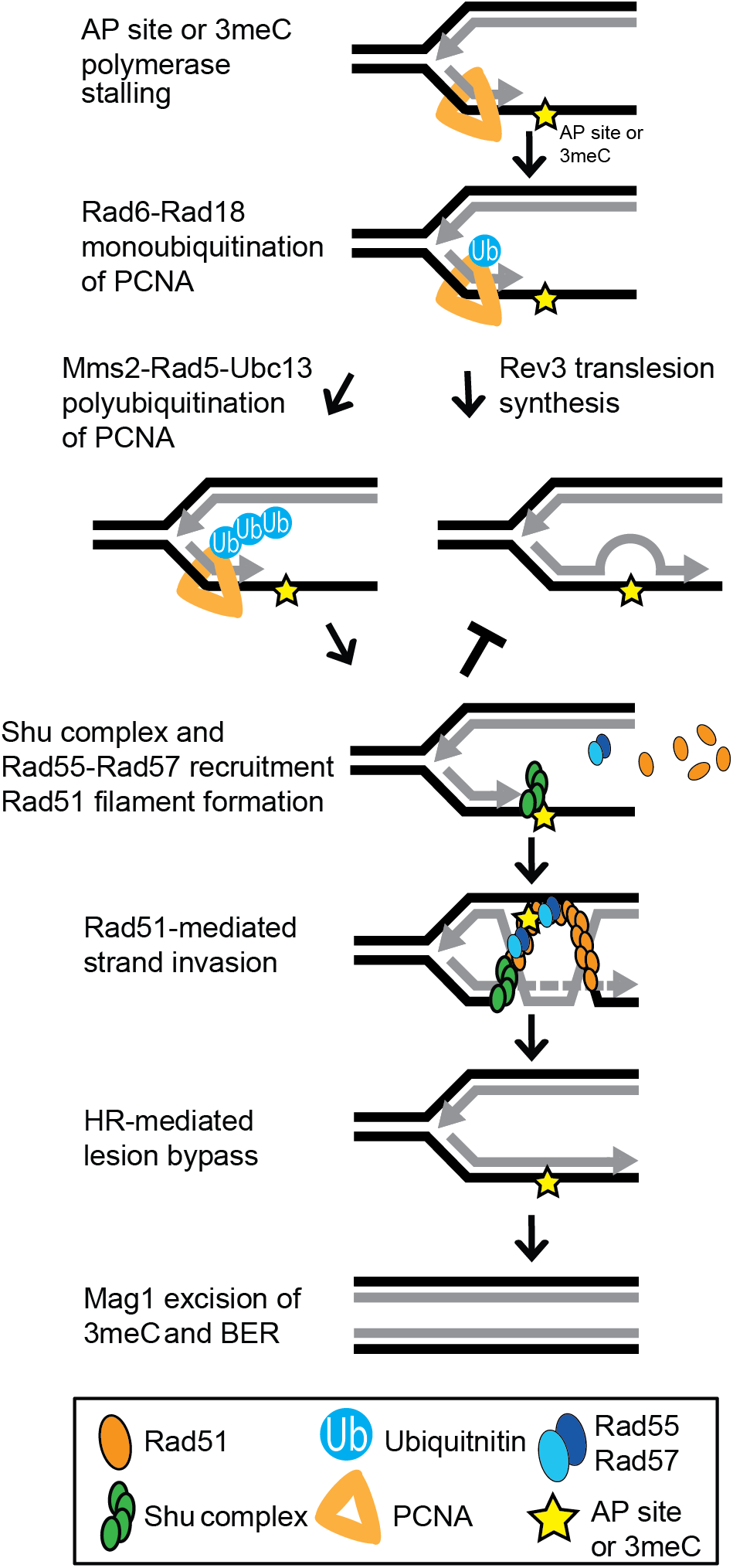
Model of Shu Complex-Mediated Error-Free Bypass of AP Sites and 3meC. MMS-induced AP sites and 3meC (yellow star) arising at DNA replication intermediates at ssDNA can stall the replicative polymerase. Replication fork stalling leads to PCNA (orange triangle) K63–linked polyubiquitination of lysine 164 (K164) by the sequential activities of the Rad6-Rad18 and Mms2-Rad5-Ubc18 complexes. The Shu complex (green ovals) through its DNA-binding components, Csm2-Psy3, binds to 3meC at a double-flap DNA junction to promote Rad55-Rad57 (blue ovals) recruitment and Rad51 filament formation (orange ovals). Thus, enabling Rad51-mediated HR with the newly synthesized sister chromatid. Importantly, the Shu complex activity prevents mutagenesis from TLS-mediated error-prone bypass of 3meC. After DNA synthesis using the undamaged sister chromatid as template, the HR intermediates are resolved. The error-free bypass of 3meC enables S-phase completion in a timely manner. Finally, after replication is completed, 3meC are likely recognized and excised by the Mag1 glycosylase, which initiates the BER-mediated repair.

It is important to note that TLS can bypass 3meA directly and can also contribute to the mutation pattern observed for both WT and *csm2*Δ cells. Direct TLS bypass of 3meA leads to a mutation pattern of elevated A->T substitutions (Shrivastav et al., 2010). Therefore, we cannot rule out the contribution of direct TLS bypass of 3meA to the mutation pattern observed in our sequence analysis. However, previous work from our group has provided genetic, *in vivo*, and *in vitro* evidence of the Shu complex role in the error-free bypass of AP sites specifically (Godin 2016, Rosenbaum 2019).

AlkB proteins are also able to repair 1meA, which primarily occurs at ssDNA (Fedeles et al., 2015). Like 3meC, 1meA is a toxic and mutagenic adduct (Shrivastav et al., 2010). It possible that this lesion is also bypassed by the Shu complex and the error-free PRR pathway. Hence, we cannot rule out that ALKBH2 repair of 1meA may also contributes to the rescue of MMS-induced phenotypes that we observe. However, it is evident from our sequence analysis that the Shu complex contributes to the bypass of lesions occurring at G:C base pairs, which excludes 1meA. Moreover, 3meC is likely the main lesion contributing to mutagenesis from MMS at ssDNA (Saini et al., 2020, Yang et al., 2010).

A previous study claimed that yeast *TPA1* is an AlKB homolog (Shivange, Kodipelli et al., 2014). However, in our hands, and consistent with a recent report (Admiraal et al., 2019), *tpa1Δ* cells show no increased sensitivity to MMS. Furthermore, unlike other genes that are involved in repairing MMS-induced lesions, we find that *TPA1* does not exhibit a synthetic sick phenotype with Shu complex mutants upon MMS exposure (**Figure S4**). Recently, the Mag1 glycosylase was shown to excise 3meC in dsDNA; therefore, initiating their repair through BER (Admiraal et al., 2019). However, Mag1 cannot excise 3meCs from ssDNA (Admiraal et al., 2019). Whole genome sequencing of MMS-treated *mag1*Δ yeast also contain primarily an elevation of substitutions at A:T base pairs indicating that Mag1 activity in yeast contributes little to protecting against 3meC-induced mutations. This is likely due to the restriction of these lesions to ssDNA and highlights the need for a mechanism to bypass this lesion during DNA replication (**Figure 6**). Mag1 likely plays a significant role in the removal of 3meC from dsDNA after its Shu complex-mediated bypass allows replication to be completed (**Figure 6**).

Previous work from our group and others show that the Shu complex role is functionally conserved in humans and mice (Abreu, Prakash et al., 2018, Martino & Bernstein, 2016, Martino, Brunette et al., 2019). Additionally, human cells containing Shu complex deletions are sensitive to MMS and DNA alkylating agents. Therefore, future studies will address whether the human Shu complex functions as a back-up of the ALKBH enzymes in tolerating mutagenesis and toxicity from 3meC. This is of particular importance since both *ALKBH2* and *ALKBH3* have been proposed as tumor suppressors, being silenced in various tumors, including gastric and breast cancer (Fedeles et al., 2015, Gao, Li et al., 2011, Knijnenburg, Wang et al., 2018, Stefansson, Hermanowicz et al., 2017). On the other hand, *ALKBH3* is often overexpressed in different cancers and inhibition of *ALKBH2* and *ALKBH3* can sensitize cancer cells to alkylating chemotherapy (Choi, Jang et al., 2011, Koike, Ueda et al., 2012, Tasaki, Shimada et al., 2011, Wang, Wu et al., 2015, Wu, Xu et al., 2011). Moreover, *ALKBH2* and *ALKBH3* upregulation mediates resistance to chemotherapeutic agents such as temozolomide and *ALKBH3* loss leads to endogenous 3meC accumulation in tumors cell lines (Cetica, Genitori et al., 2009, Dango et al., 2011, Johannessen, Prestegarden et al., 2013). A connection between the human RAD51 paralog, RAD51C, and ALKBH3 was recently described where RAD51C was important for recruitment of ALKBH3 to DNA with 3meC. A role of the Shu complex promoting tolerance of 3meC could provide a new avenue for therapeutic approaches to target these tumors.

## MATERIALS AND METHODS

### Yeast strains, plasmids, and oligos

The strains utilized are listed in **Supplemental Table 1** whereas all oligonucleotides used are listed in **Supplemental Table 2**. The Y2H strains PJ69-4A and PJ69-4a were used as described (Godin et al., 2013, James, Halladay et al., 1996). All strains are isogenic with W303 RAD5+ W1588-4C (Thomas & Rothstein, 1989) and W5059-1B (Zhao, Muller et al., 1998). KBY-1088-3C (rad55-S2,8,14A) was generated by transformation of a cassette containing the 50 bp homology upstream of the *RAD55* start codon and the rad55-S2,8,14A ORF fused to a kanMX6 resistance cassette and the 50 bp homology downstream of the *RAD55* stop codon. This fused cassette was obtained using Gibson Assembly^®^ Master Mix (NEB) following the manufacturer’s instructions and the primers used to generate the assembly fragments were designed using NEBuilder Assembly Tool (https://nebuilder.neb.com). The rad55-S2,8,14A gene fragment was commercially synthesized whereas the kanMX6 cassette was amplified from the pFA6a-kanMX6 plasmid (Longtine, McKenzie et al., 1998). Prior to transformation, the fused product was PCR amplified using primers with 50nt of homology to the flanking regions of *RAD55* as described (Longtine et al., 1998). All yeast transformations were performed as described (Sherman, Fink et al., 1986). The integration of the rad55-S2,8,14A ORF was verified by PR amplification followed by sequencing using the KanHisNat and Rad55.Check oligos as described in as described (Longtine et al., 1998). pGAD-rad55-S2,8,14A was generated using Gibson Assembly^®^ Master Mix (NEB) following the manufacturer’s instructions and the primers used to generate the assembly fragments were designed using NEBuilder Assembly Tool (https://nebuilder.neb.com). ALKBH2 and ALKBH3 were cloned in the pAG416GPD-ccdB vector (Plasmid #14148 Addgene). pAG416GPD-ccdB-ALKBH2-CD was generated by site-directed mutagenesis of the pAG416GPD-ccdB-ALKBH2 plasmid as described (Zheng, Baumann et al., 2004) with minor adaptations according to the manufacturer’s recommendations for PCR using Phusion High-Fidelity PCR Master Mix with HF Buffer (Thermo). All knock-outs were generated using the S1 and S2 primers and knock-out cassettes as described in (Longtine et al., 1998). All plasmids and strains were verified by DNA sequencing.

### Chronic MMS Exposure and DNA Whole Genome Sequencing

Individual colonies of WT or *csm2Δ* cells were grown overnight at 30°C. The cultures were then pinned onto YPD medium containing 0.008% MMS using a pinning robot from S&P Robotics. After a 2-day incubation at 30°C the plates were replica-plated onto YPD plates containing 0.008% MMS using a robotic pinner, and then replated onto fresh YPD medium containing 0.008% MMS a total of 10 times. The MMS-exposed yeast were separated into single colonies (96 per strain). These colonies were inoculated in YPD cultures and grown overnight at 30°C. Genomic DNA was extracted from each culture by resuspending the yeast pellets in lysis buffer (20 mM Tris-Cl, 200 mM LiAc, 1.5% SDS, pH7.4), incubating the yeast for 15 min at 70°C, incubating on ice for 10 min, adding an equal volume of 4M NaCl, and centrifuging the samples at maximum speed for 30 minutes at 4°C. The supernatant was then added to an equal volume of phenol:chloroform:isoamyl alcohol (PCI). An equal volume of isopropanol was added to the aqueous phase of the PCI extraction. The DNA pellet was washed twice with 70% ethanol prior to resuspending in 10 mM Tris 8 buffer. The DNA the subjected to Illumina whole genome sequencing. 1 μg of genomic DNA per sample was sheared and used to generate libraries for sequencing with a KAPA DNA HyperPrep kit. Multiplexed libraries were sequenced with 96 samples on a single lane of an Illumina Hiseq4000. Illumina reads were aligned to the Saccer3 S288C reference genome with CLC Genomics Workbench version 7.5. CLC Genomics Workbench was also used to identify mutations from these alignments using methods similar to those previously described (Sakofsky, Roberts et al., 2014). Briefly, mutations were identified as differences from the reference genome that were greater than 9 reads covering the site, and for which between 45-55% of the reads supported the mutation. Additionally, any mutations occurring in multiple samples were removed from analyses as likely polymorphisms or alignment artifacts. The complete list of mutations is provided in **Supplemental Table 3**. Raw sequencing reads in fastq format have been submitted to the NCBI short read archive under BioProject accession number PRJNA694993.

### Growth Assays

Individual colonies of the indicated strains were transformed with an empty plasmid or a plasmid expressing ALKBH2 as indicated. The cultures were grown in 3ml YPD or SC-URA medium overnight at 30°C. Five-fold serial dilutions were performed (Godin et al., 2016c) except that 5 μL of culture at 0.2 OD_600_ were 5-fold serially diluted onto YPD medium or YPD medium containing the indicated MMS concentration. UV treatment was performed using Stratagene Stratalinker 2400 UV Crosslinker. The plates were imaged after 48 h or 72 h of incubation at 30°C and the brightness and contrast was globally adjusted using Photoshop (Adobe Systems Incorporated).

### Protein Blotting

Parental and *csm2*Δ cells were transformed with an empty plasmid or plasmid expressing either ALKBH2 or ALKBH2-CD. The cultures were grown in SC-URA medium overnight at 30°C. Subsequently, equal cell numbers (1 ml 0.75 OD_600_) from each culture were pelleted, supernatant was removed, and washed once with ddH_2_O and pelleted again. Protein was extracted from whole cell lysates by TCA preparation as described in 51 μl of loading buffer (Knop et al. 1999). The 13 μl of protein was run on a 10% SDS-PAGE gel and transferred to a PDVF membrane by semidry transfer (Bio-Rad) at 13V for 90 min. ALKBH2 or Kar2 was Western blotted using αALKBH2 antibody [AbCam (ab154859); 1:1000 with secondary antibody anti-rabbit (Jackson ImmunoResearch Laboratories 1:10,000] or αKar2 antibody [Santa Cruz (sc-33630); 1:5000 with secondary antibody anti-rabbit (Jackson ImmunoResearch Laboratories 1:10,000] as a loading control.

### Survival Assays

Individual colonies of WT or *csm2Δ* cells were transformed with an empty plasmid or a plasmid expressing *ALKBH2* and grown in 3ml SC-URA medium at 30°C overnight. The cultures were diluted to 0.2 OD_600_ in 50 ml SC-URA medium and grown for 3-4 h at 30°C. The cultures were all diluted to 0.2 OD_600_ in YPD or YPD containing 0.05%, 0.1% or 0.2% MMS and incubated for 30 minutes at 30°C. After the treatment, the cultures were washed twice with YPD and resuspended to 0.2 OD_600_ in YPD. The cultures were diluted 1/10000 (untreated and 0.05% MMS) or 1/1000 (0.1% and 0.2% MMS) and 150 ul were plated in YPD medium plates in duplicate. The plates were incubated at 30°C for 2 days before imaging. The colonies were counted using OpenCFU (Geissmann, 2013). Data from 5 to 9 colonies from at least 3 independent transformants was used.

### Canavanine Mutagenesis Assay

Individual colonies of the indicated strains were transformed with an empty plasmid or a plasmid expressing *ALKBH2* and grown in 3ml SC-URA medium or SC-URA medium containing 0.00033% MMS for 20 h at 30°C. The cultures were diluted to 3.0 OD_600_. 150 ul were plated on SC-ARG+CAN (0.006% canavanine) medium in duplicate or 150 ul of a 1:10,000 dilution were plated on SC medium in duplicate. The plates were then incubated for 48 hours at 30°C before imaging. The colonies were counted using OpenCFU (Geissmann, 2013) to measure total cell number (SC) or forward mutation rates (SC-ARG+CAN). The mutation frequency was obtained by dividing colony number in SC-ARG+CAN by the number obtained in the SC plates times the dilution factor. Mutation frequencies were converted to mutation rates using methods described (Drake, 1991). Data from 10 to 20 colonies from at least three independent transformants was used. Differences in mutation rates were evaluated using a Mann-Whitney rank sum test comparing the independent rate measurements between untreated *csm2*Δ yeast expressing either an empty vector control or *ALKBH2* and between MMS-treated *csm2*Δ yeast expressing either an empty vector control or *ALKBH2*. Mutations occurring within the *CAN1* gene were identified by selecting independent canavanine resistant colonies, isolating genomic DNA from these samples, and PCR amplifying the *CAN1* gene from each isolate with primers containing unique barcodes (**Supplemental Table 4**). These PCR products were pooled and sequenced on a PacBio Sequel sequencer as in (Hoopes et al., 2017). PacBio reads were separated by barcodes and aligned to the *CAN1* gene sequence using Geneious software. Geneious was also used to identify mutations among these alignments as differences from the reference *CAN1* sequence that are supported by two or more independent reads that occur in greater than 30% of all reads for each sample. 452 out of 458 mutations identified were supported by greater than 50% of the reads for the sample. All mutations identified by PacBio sequencing are provided in **Supplemental Table 5**.

### Yeast-Two-Hybrid Assays

The yeast-two-hybrid experiments using the indicated pGAD and pGBD plasmids were performed (Godin et al., 2013) except that both pGAD and pGBD plasmids were transformed into PJ69-4A (James et al., 1996). A yeast-two-hybrid interaction is indicated by growth on synthetic complete (SC) medium lacking histidine, tryptophan, and leucine whereas equal cell loading was observed by plating the cells on SC medium lacking tryptophan and leucine to select for the pGAD (leucine) or pGBD (tryptophan) plasmids.

### Csm2-Psy3 Purification

The Csm2-Psy3 heterodimer was cloned into the dual expression plasmid pRSFDuet (EMD Millipore) which encode a Strep-tagged Csm2 and FLAG-tagged Psy3. All affinity tags were fused to the N-terminus of the protein. This plasmid was transformed into *E.coli* (Rosetta P.LysS) and grown at 37°C until 0.6 OD_600_ and recombinant protein expression was induced by addition of 0.5 mM isopropyl beta thiogalactoside (IPTG) at 18°C overnight for 16–18 h. Cells were harvested by centrifugation and pellets were frozen at −80°C. Approximately 38 g of cell pellet was lysed in 80 mL of lysis buffer containing 25 mM Tris (pH 7.4), 300 mM KCl, 10% glycerol, 5 mM β-mercaptoethanol supplemented with protease inhibitors (Roche), 1mM PMSF, 1 mM ZnCl2 and 0.01% IGEPAL CA-630. Cells were lysed using an emulsiflex and centrifuged at 15,000 × *g* for 1 h at 4°C. Lysed supernatant was incubated with 5 mL of ANTI-FLAG M2 affinity resin (Sigma) overnight and then washed in a solution consisting of 25 mM Tris (pH 7.4), 150 mM KCl, 10% glycerol, and 0.01% IGEPAL CA-630. To elute bound Csm2-Psy3, the wash buffer was supplemented with 0.1mg/mL 3X-FLAG peptide (Sigma). Elutions were then loaded onto a HiTrap Heparin HP (GE Healthcare) affinity chromatography. The complex was loaded onto the column and washed for 200 mL with (25 mM Tris (pH 7.4), 150 mM KCl, 1 mM DTT, 10% glycerol, and 0.01% IGEPAL CA-630). The complex was eluted with a gradient elution from 0% to 100% (25 mM Tris (pH 7.4), 150 mM KCl, 1 mM DTT, 10% glycerol, and 0.01% IGEPAL CA-630) over 20 mL. The Csm2-Psy3 protein typically eluted around 400–600 mM NaCl. Then, Csm2-Psy3 Heparin elutions were subsequently purified by size exclusion chromatography using a Sephacryl S200 column (GE Healthcare) in buffer (25 mM Tris (pH 7.4), 150 mM KCl, 1 mM DTT, 10% glycerol, and 0.01% IGEPAL CA-630) eluting as a single peak and visualized as heterodimers by SDS-PAGE electrophoresis. Csm2-Psy3 protein concentration was determined by BCA Assay as previously described (Rosenbaum et al., 2019).

### Substrates for Fluorescence Anisotropy

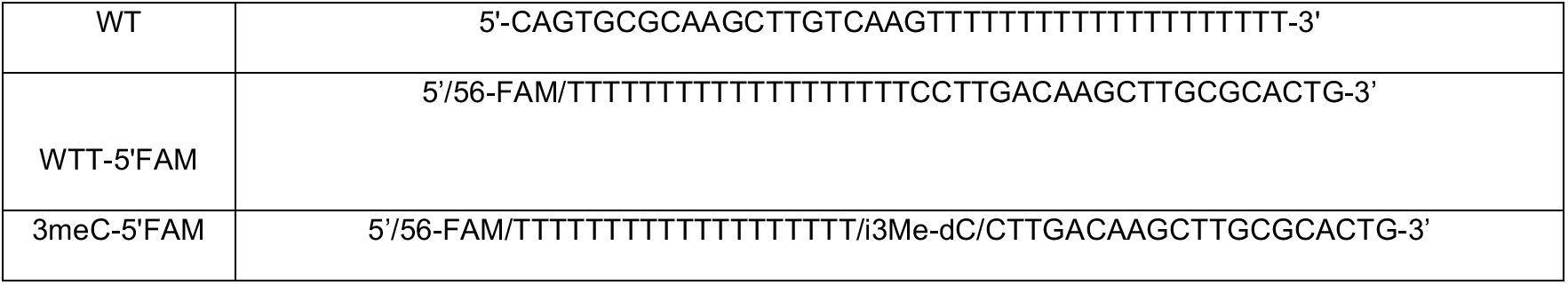

### Equilibrium Fluorescence Anisotropy Assays

Anisotropy experiments were performed using a FluoroMax-3 spectrofluorometer (HORIBA Scientific) on and a Cary Eclipse Spectrophotometer. All substrates used for anisotropy contained a 5’-FAM moity incorporated at the end of the single-stranded end of the fork (**Figure 4C**). Substrates either contained a thymidine (T) or 3meC at the double-flap junction in the ssDNA. Anisotropy measurements were recorded in a 150 μL cuvette containing 20 mM Tris pH 8.0 and 5.11 or 4.53 nM of FAM-labeled double-flap substrate as a premixed sample of purified Csm2-Psy3 protein and substrate (5.11 nM WT +1.6 μM Csm2-Psy3 dimer in 50 nM NaCl) was titrated into the cuvette. Fluorescence anisotropy measurements were recorded using the integrated polarizer and excitation and emission wavelengths of 495 and 520 nm, respectively, with slit widths of 10 nm. Titrations were carried out at 25°C and were carried out until anisotropy became unchanged. All experiments were performed in triplicate. Dissociation constants (*K*_d_) were calculated by fitting our data to a quadratic equation [*Y*=*M*\* ((*x* + *D* + *K*_d_)–sqrt(((*x* + *D* + *K*_d_)^2^)–(4\**D*\**x*)))/(2\**D*)] assuming a one site binding model. Data were fit with PRISM7 software. Unprocessed raw anisotropy values are in the source data file.

## ACKNOWLEDGEMENTS

This study was supported by the National Institutes of Health grant (ES030335 to K.A.B.; CA218112 to S.A.R.) and the American Cancer Society (129182-RSG-16-043-01-DMC to K.A.B. and 133947-PF-19-132-01-DMC to S.R.H.).

## SUPPLEMENTAL FIGURES

**Supplemental Figure 1.**
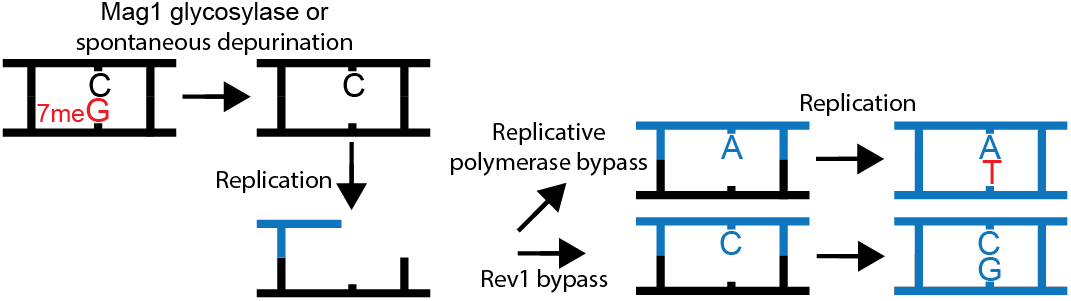
Schematic of How 7meG Derived AP Sites Lead to G to T as the Main Substitution Pattern. 7meG can be converted to AP sites by spontaneous hydrolysis or the glycosylase activity of Mag1. AP may be bypassed by the TLS activity of Rev1 which is this case is error-free. Alternatively, lesion bypass by the replicative polymerase, which predominantly incorporates A across AP sites, would result in a G to T mutation pattern.

**Supplemental Figure 2.**
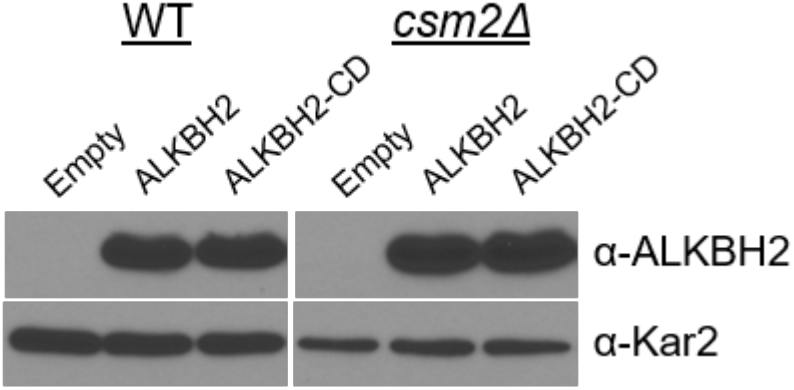
Protein Blot Analysis of ALKBH2 and ALKBH2-CD Expression in WT and *csm2*Δ Cells. WT and *csm2*Δ strains expressing an empty plasmid (pAG416GPD-ccdB) or a plasmid (pAG416GPD-ccdB) expressing either ALKBH2 or ALKBH2-CD were analyzed for ALKBH2 protein levels. Protein extracts from equal cell numbers were analyzed by western blot for ALKBH2 (α-ALKBH2) or Kar2 (α-Kar2) expression as a loading control.

**Supplemental Figure 3.**
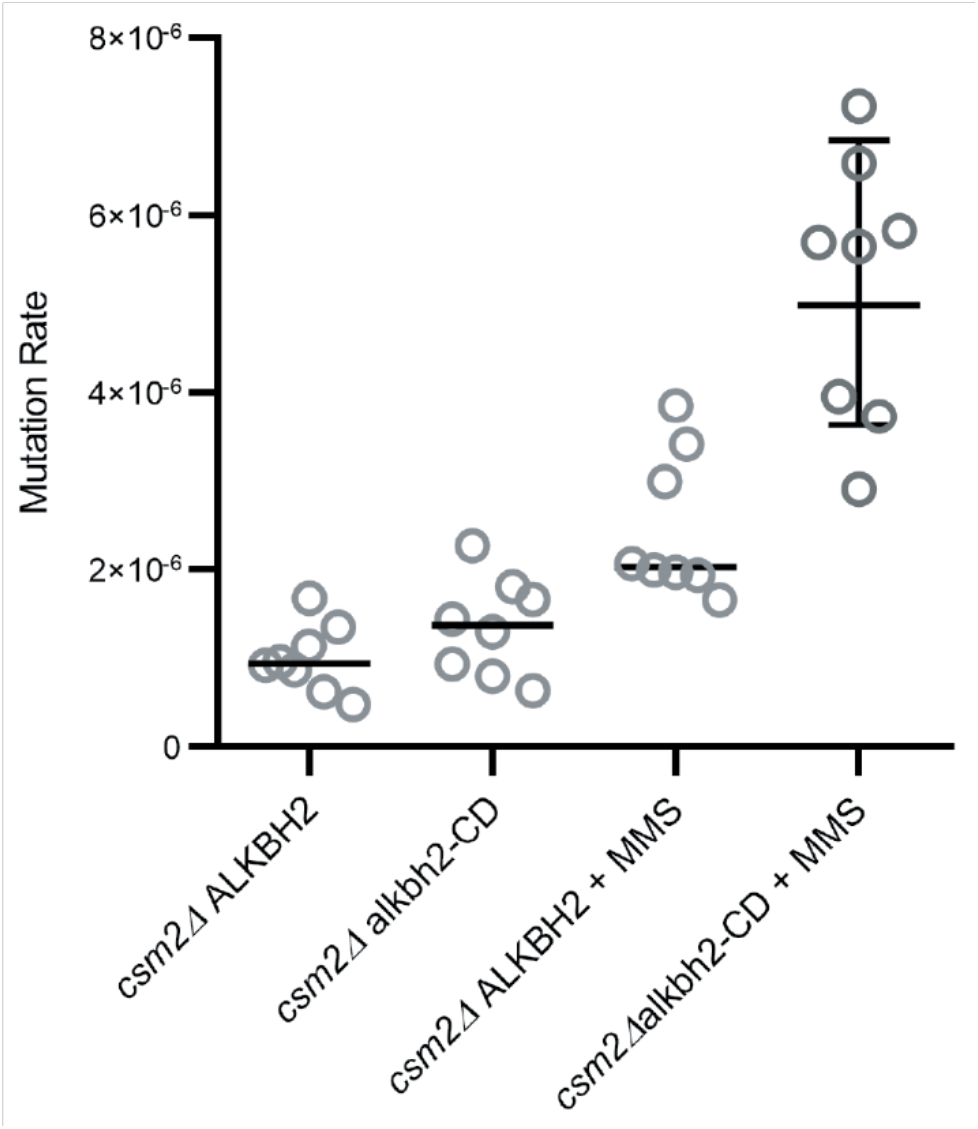
Expression of a *ALKBH2* Catalytic Dead Mutant Does Not Rescue the Mutagenesis Observed in MMS Exposed *csm2Δ* Cells. *csm2Δ* cells expressing *ALKBH2-CD* exhibit an increased MMS-induced mutation rate. Spontaneous and MMS-induced mutation rates at the *CAN1* locus were measured in *csm2*Δ cells transformed with either a plasmid expressing *ALKBH2* or *ALKBH2-CD*. Each measurement (dots) and the median value of 8 experiments (horizontal bar) were plotted. The *p*-values between *ALKBH2* and *ALKBH2-CD* expressing cells were calculated using a Mann-Whitney Ranked sum test and were p > 0.05 and p < 0.03 for the for the untreated and MMS treated samples, respectively.

**Supplemental Figure 4.**
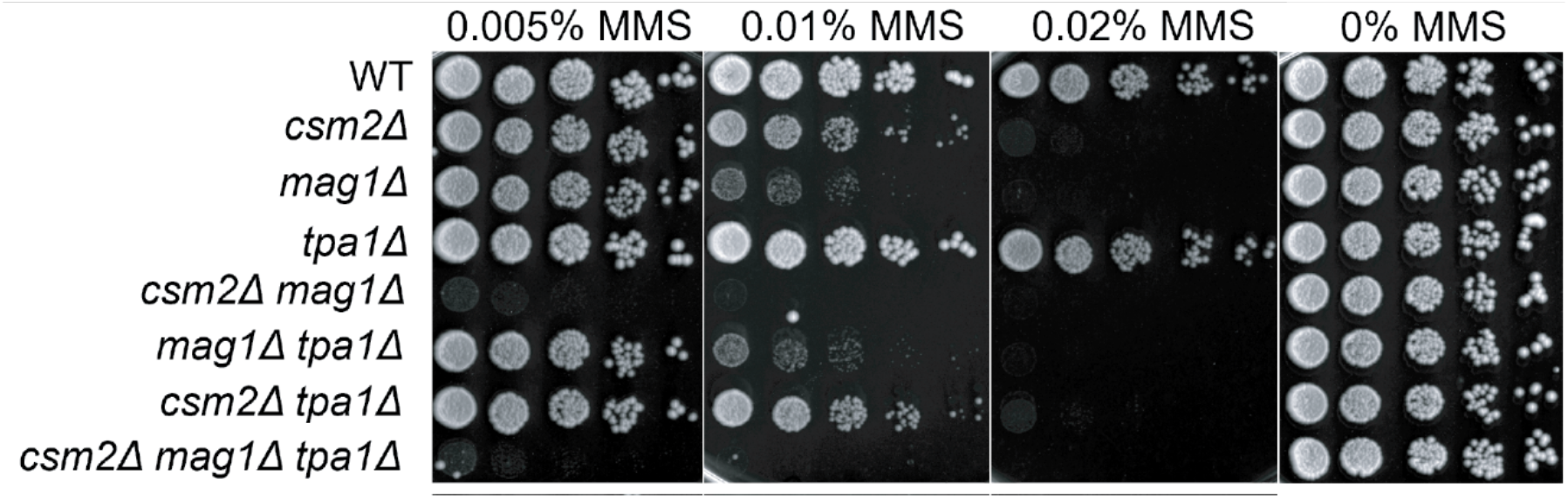
*TPA1* Does Not Genetically Interact with *CSM2* or *MAG1* for MMS Damage. Five-fold serial dilution of cells with the indicated genotypes were transformed onto rich YPD medium or rich YPD medium containing the indicated MMS concentration were incubated for two days at 30°C prior to being photographed.

